# Maximizing the reliability and the number of species assignments in metabarcoding studies

**DOI:** 10.1101/2022.11.17.513905

**Authors:** Audrey Bourret, Claude Nozères, Eric Parent, Geneviève J. Parent

## Abstract

The use of environmental DNA (eDNA) for biodiversity assessments has increased rapidly over the last decade. However, the reliability of taxonomic assignments in metabarcoding studies is variable, and affected by the reference databases and the assignment methods used. Species level assignments are usually considered as reliable using regional libraries but unreliable using public repositories. In this study, we aimed to test this assumption for metazoan species detected in the Gulf of St. Lawrence, in the Northwest Atlantic. We first created a regional library with COI barcode sequences including a reliability ranking system for species assignments. We then estimated the accuracy of the public repository NCBI-nt for species assignments using sequences from the regional library, and contrasted assigned species and their reliability using NCBI-nt or the regional library with a metabarcoding dataset and popular assignment methods. With NCBI-nt and sequences from the regional library, Blast-LCA was the most accurate method for species assignments but the proportions of accurate species assignments were higher with Blast-TopHit (>80 % overall taxa, between 70 and 90 % amongst taxonomic groups). With the metabarcoding dataset, the reliability of species assignments was greater using the GSL-rl compared to NCBI-nt. However, we also observed that the total number of reliable species assignments could be maximized using both GSL-rl and NCBI-nt, and their optimal assignment methods, which differed. The use of a two-step approach in species assignments, using a regional library and a public repository, could improve the reliability and the number of detected species in metabarcoding studies.

## Introduction

The use of environmental DNA (eDNA) for biodiversity assessments and monitoring has increased rapidly over the last decade given the high potential of this non-intrusive approach to uncover biodiversity with limited effort (Taberlet et al. 2012, Makiola et al. 2020). eDNA metabarcoding surveys collect and detect traces of a diversity of organisms in various types of environmental samples using high-throughput sequencing or PCR-based approaches (Taberlet et al. 2012, Yu et al. 2012). Surveys of eDNA generally involve a series of steps such as sample collection, extraction, targeted amplification, high-throughput sequencing, and bioinformatic processing, which includes taxonomic assignments to reference sequences from a public repository or a regional library (Deiner et al. 2017). Only a small fraction of detected eDNA sequences in environmental samples can currently be assigned to a species-level identity owing to a lack of data and taxonomic resolution in publicly available resources (Deiner et al. 2017, Leite et al. 2021, Zafeiropoulos et al. 2021). The reliability and precision of taxonomic assignments is affected by the quality and availability of sequences in repositories and the assignment methods, thereby limiting confidence in the use of eDNA for biodiversity monitoring and targeted species detections (Coissac et al. 2012, McGee et al. 2019, Meiklejohn et al. 2019, Gold et al. 2021, Hleap et al. 2021).

Several public repositories exist and can be used as reference databases to provide taxonomic assignments in metabarcoding studies. The public National Center for Biotechnology Information Nucleotide database (NCBI-nt, including the well-known GenBank database) is the largest sequence repository and is widely used in eDNA metabarcoding studies (Porter and Hajibabaei 2018b, 2020). However, the presence of mislabeled specimens, the large variation in quality of sequences available, and gaps in species coverage (i.e., unrepresented species) result in erroneous species identification when directly comparing unknown sequences to NBCI-nt (Bidartondo 2008, Mioduchowska et al. 2018, Leray et al. 2019). The Barcode of Life Data Systems (BOLD) is another sequence repository specific to the most common barcode regions, including the cytochrome c oxidase I (COI) gene which is the widely used gene region for animal DNA barcoding (Ratnasingham and Hebert 2007, Porter and Hajibabaei 2018b). BOLD displays mandatory (e.g., institution storing voucher specimen, sampling country) and optional (e.g., sampling location, specimen photos) metadata, performs groupings of similar sequences into Barcode Index Number (BIN), and permits editing or updating of records, all of which assists with data quality control. However, like GenBank, it is also vulnerable to submissions of misidentified specimens (McCusker et al. 2013, Oliveira et al. 2016, Fontes et al. 2021, Radulovici et al. 2021). As reliable taxonomic assignments at the species-level are expected under many regulatory contexts (e.g., environmental status assessment, monitoring of invasive species or species at risks; Aylagas et al. 2014, Hering et al. 2018, Bush et al. 2019, Piper et al. 2019), some metabarcoding studies have questioned the value of using public repositories (e.g., von Ammon et al. 2018, Locatelli et al. 2020, Gold et al. 2021). Characterizing the proportion of accurate species assignments using NCBI-nt would be highly valuable to understand the extent of uncertainty in species eDNA detection and consequently, enable an accurate interpretation of a metabarcoding study’s results.

Alternatively, curated regional libraries have been shown to reduce errors in species assignments (Gold et al. 2021). Regional libraries are limited to species expected in predefined areas, and can be created by data mining and curating existing sequences from public repositories and/or from generating sequences from specimens. They have the advantage to limit spurious assignments to related but non-local species, and to reveal gaps (i.e., missing sequences) in taxonomic groups (Weigand et al. 2019, Ramirez et al. 2020, Jazdzewska et al. 2021). Examples of regional libraries are widely available in the northern hemisphere for multiple taxonomic groups (e.g., Knebelsberger et al. 2014, Hänfling et al. 2016, Stoeckle et al. 2017, Fraija-Fernández et al. 2020, Gold et al. 2021, Van Den Bulcke et al. 2021). Some of these reference libraries present ranking systems to ensure high taxonomic reliability (e.g., Costa et al. 2012, Knebelsberger et al. 2014). Ranking systems are often provided to target future barcoding efforts and improvements in reference sequences. No explicit ranking system about the uncertainty of species assignment has yet been presented within metabarcoding studies. Such a system would be highly valuable to provide clear indications on the reliability of species assignments for eDNA end-users.

Another source of variability in species assignments are the bioinformatics software and pipelines used in metabarcoding studies. Recently, studies have started to evaluate the accuracy of taxonomic assignments using various bioinformatic methods (O’Rourke et al. 2020, Hleap et al. 2021, Mathon et al. 2021). These studies compared taxonomic assignment methods that are based on strategies such as alignment, composition, or modelling (Richardson et al. 2017, see also four strategies in Hleap et al. 2021). The Basic Local Alignment Search Tool (BLAST) is an alignment-based approach extensively used in metabarcoding studies that relies on a nearest-neighbor approach to return best hits between unknown sequences and records from a reference database (Camacho et al. 2009). The taxonomic identity of the unknown sequence may be inferred in conjunction with a least common ancestor (LCA) or a Top Hit approach with identity threshold, usually between 95 and 99%. These thresholds should reflect expected inter-species divergence, but high variation among taxonomic groups may cause pitfalls in assignments (Wang et al. 2007, Alberdi et al. 2018). Classifiers using machine-learning algorithms are from composition-based approaches that have shown good performance in some contexts of species assignments (Richardson et al. 2017, Murali et al. 2018, Porter and Hajibabaei 2018a). They can take advantage of phylogeny between reference sequences and thus are less affected by shared divergence between groups. Classifiers are trained on a reference library, and pre-trained classifiers are increasingly available (e.g., Porter and Hajibabaei 2018a). However, recent benchmarking studies have shown lower performance of classifiers compared to BLAST (O’Rourke et al. 2020, Hleap et al. 2021, Mathon et al. 2021).

This study aimed to estimate the accuracy of species assignments using NCBI-nt and to contrast the reliability of using NCBI-nt or a regional library on a metabarcoding dataset with popular assignment methods (Fig. 1). To achieve these objectives, we first created a curated regional library (GSL-rl) using publicly available sequences from BOLD for COI barcode locus of metazoans from the Gulf of St. Lawrence. The regional library presents a reliability ranking system for species assignments based on sequence availability and similarity that can be understood by any eDNA end-users, scientists or not. We then used sequences from GSL-rl to estimate the accuracy of NCBI-nt. We also compared the assigned species in a metabarcoding dataset using NCBI-nt or GSL-rl and their reliability. We reached the conclusion that using a two-step approach, i.e., species assignments with a regional library and a public repository, is desirable to maximize the reliability and the number of species assignments in metabarcoding studies.

**Figure 1.**
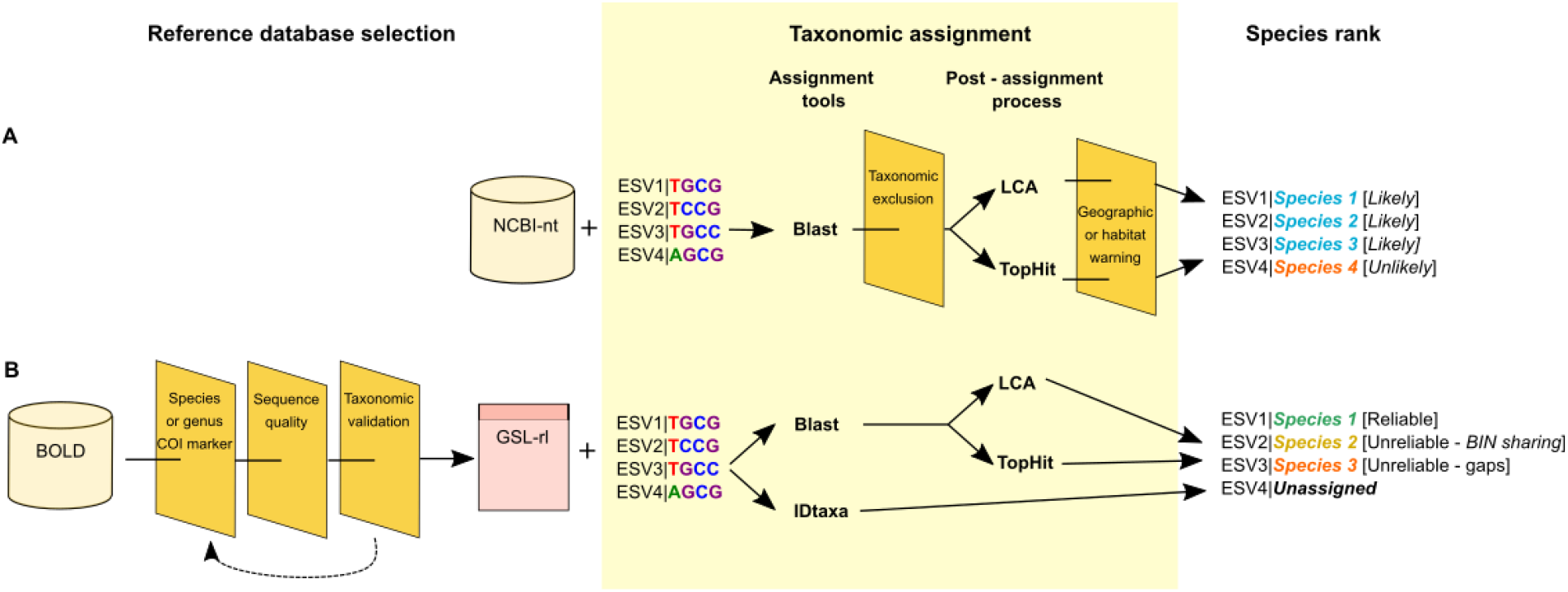
Schematic representation of the two sources of reference sequences used and compared in this study and their associated methods, A) the public repository NCBI nucleotide database (NCBI-nt), and B) the newly created regional library for metazoans from the Gulf of St. Lawrence (GSL-rl). Using NCBI-nt, exact sequence variants (ESVs) were assigned using the BlastX tool *blastn* (hereafter Blast; Camacho et al. 2009). Assignment results were filtered based on taxonomic identity, then a least common ancestor (LCA) or a TopHit method was used to assign a unique taxon identity to each ESV. The creation of GSL-rl involved data mining of the public repository BOLD and included multiple filtering and auditing steps and a feedback loop to improve it. Taxonomic assignments of ESV were performed with Blast or with the classifier IDtaxa (Murali et al. 2018). For GSL-rl, the species ranking was based on sequence availability and sequence similarity to closely related species in the Gulf of St. Lawrence. For NCBI-nt, the species ranking involved a plausibility filter based on the location (see methods for more details).

## Methods

### Creation of a regional library for the Gulf of St. Lawrence (GSL-rl) with a reliability ranking system

The creation of a curated regional library for the Gulf of St. Lawrence was composed of three major steps: 1) obtaining a list of marine faunal species from both decision-makers and available regional taxonomical information (Nozères 2017), 2) creating an initial regional library using bioinformatics tools with BOLD and rounds of revisions based on quality and similarity of sequences, and 3) enhancing the draft regional library using a metabarcoding dataset (hereafter GSL regional library: GSL-rl; Fig. S1). All sequences retained in the GSL-rl had names at the genus or species level and were already published on BOLD. More details about the creation of the GSL-rl are provided in the supplementary material.

We created the GSL-rl to identify molecular operational taxonomic units (MOTUs) at the species level. Each species in the GSL-rl was ranked based on sequence availability and similarity (Table S1). Species with reference sequences for itself and closely related species (i.e., from the same genus) acknowledged to be present in the Gulf of St. Lawrence were ranked as “Reliable” if they did not share BOLD’s barcode index number (BIN; i.e., a unique identifier of sequences based on genetic distance, Ratnasingham and Hebert 2013). Species with reference sequences for itself, but not for all congeners acknowledged to be present in the Gulf of St. Lawrence (i.e., other species of the same genus) were ranked as “Unreliable due to gaps”. Species with reference sequences sharing BIN with other species were ranked as “Unreliable due to BIN sharing”. Common causes of BIN sharing are genetic similarities between species or specimen misidentification. For the GSL-rl, curation and validation process done during its creation should limit the BIN sharing due to specimen misidentification. Taxonomic assignments belonging to one of the two “Unreliable” categories should be interpreted with caution, and preferably not at the species-level.

### Evaluating the accuracy of species assignments using the public NCBI-nt repository

We used the curated sequences from the GSL-rl to estimate three performance parameters using NCBI-nt: 1) the proportion of assignments, 2) the proportion of accurate assignments, and 3) the accuracy.

Taxonomic assignments were performed using NCBI-nt (downloaded 2020-10-23) and the BlastX tool *blastn* (v2.10.1, Camacho et al. 2009) combined with the least common ancestor (LCA; hereafter Blast-LCA) or the Top Hit methods (hereafter, Blast-TopHit) at three identity thresholds (95, 97 and 99%) from an in-house R script. The LCA method assigns the higher taxonomic rank shared by all hits above the identity threshold while the Top Hit method only used the hits with the highest probability. We excluded hits containing “environmental sample”, “uncultured” or “predicted” in their description.

We assessed the proportion of assignments at the species level, and we considered that an assignment was accurate if it matched the species identity associated in the GSL-rl. We measured accuracy as the proportion of accurate assignments over all assignments at the species level.

### Contrasting species assignments using the regional library or the public NBCI-nt repository, and popular assignment methods

We compared the detection results from an eDNA metabarcoding dataset using GSL-rl and NCBI-nt and three assignment methods (Fig. 1). The eDNA metabarcoding dataset was obtained from the analysis of water samples collected from scientific surveys in 2018 in coastal areas of the GSL, both at the surface and bottom of the water column (see supplementary material for details on the field, laboratory and bioinformatics works underlying the eDNA metabarcoding dataset). The three assignment methods were Blast-LCA, Blast-TopHit and the IDtaxa (Murali et al. 2018). Blast assignment methods were used as described in the previous section with both GSL-rl and NCBI-nt. NCBI-nt Blast results were filtered to retain only metazoan detections and remove non-marine taxa (i.e., *Homo sapiens*, Arachnida, Insecta). IDtaxa is a classifier implemented within the DECIPHER R package (Wright 2016), and was trained only with the GSL-rl. IDtaxa classifier was selected since it would be less prone to “over classification”, i.e., classification to an erroneous group when the real group is absent from the training set, compared to the popular Ribosomal Database Project (RDP) classifier (Murali et al. 2018). Taxonomic assignments with IDtaxa were obtained at three confidence thresholds (i.e., weighted fraction of bootstrap replicates assigned to a given taxa) representing moderate confidence (40%), high confidence (50%), and very high confidence (60%) in species assignments (Murali et al. 2018).

We contrasted results obtained using GSL-rl and NCBI-nt with distinct ranking systems. Species detected with the GSL-rl were classified according to the three categories of the reliability ranking system previously created: “Reliable”, “Unreliable due to gaps”, “Unreliable due to BIN sharing” (Fig. 1B, Table S2). For species assignments with NCBI-nt, we used geographic and habitat filters to classify them as “Likely” if they were part of the Gulf of St. Lawrence checklist (Nozères 2017) or present in the areas based on the World Register of Marine Species (WoRMS, WoRMS Editorial Board 2020), and “Unlikely” if not (Fig. 1A). Such filters are often applied in metabarcoding studies but the source of information for the likeliness of a species to be present is often obscure.

### Data and R scripts availability

All data and R scripts used for the creation of the GSL-rl are provided on Github: https://github.com/GenomicsMLI-DFO/GSL_COI_ref_library

## Results

### A COI regional library with a reliability ranking system for metazoans from the Gulf of St. Lawrence (GSL-rl)

The first version of the GSL-rl comprised 1304 sequences covering 439 species (158 species of vertebrates from phylum Chordata from 68 families; 281 species of invertebrates from 129 families and 9 phyla) and 11 other taxa at the genus level only (Vertebrates: 3 genera from 2 families and phylum Chordata; Invertebrates: 8 genera from 8 families and 4 phyla; Fig 2). It represented 67.4% of the taxa on the target list (651 species) used and improved during the GSL-rl creation (Vertebrates: 94.6%; Invertebrates: 58.1%; Table S1). The sequences were retrieved mostly from the Northwest Atlantic Ocean (67.8%). A total of 525 BINs were represented (Vertebrates: 159; Invertebrates: 366), with 16 BINs that were shared by at least two taxa (Vertebrates: 8; Invertebrates: 8; Table S2), and 58 taxa occupied more than one BIN (6 vertebrates with up to 3 BINs; 52 invertebrates with up to 7 BINs; Table S3). Median sequence length was 658 pb (range: 640 – 664 pb) while the mean (± sd) of missing values (N’s) was 0.002 ± 0.034 % (max < 1 %). Genetic distances were on average 0.005 (range: 0.000–0.023) within BIN and 0.122 (range: 0.012–0.347) between intraspecific BINs.

**Figure 2.**
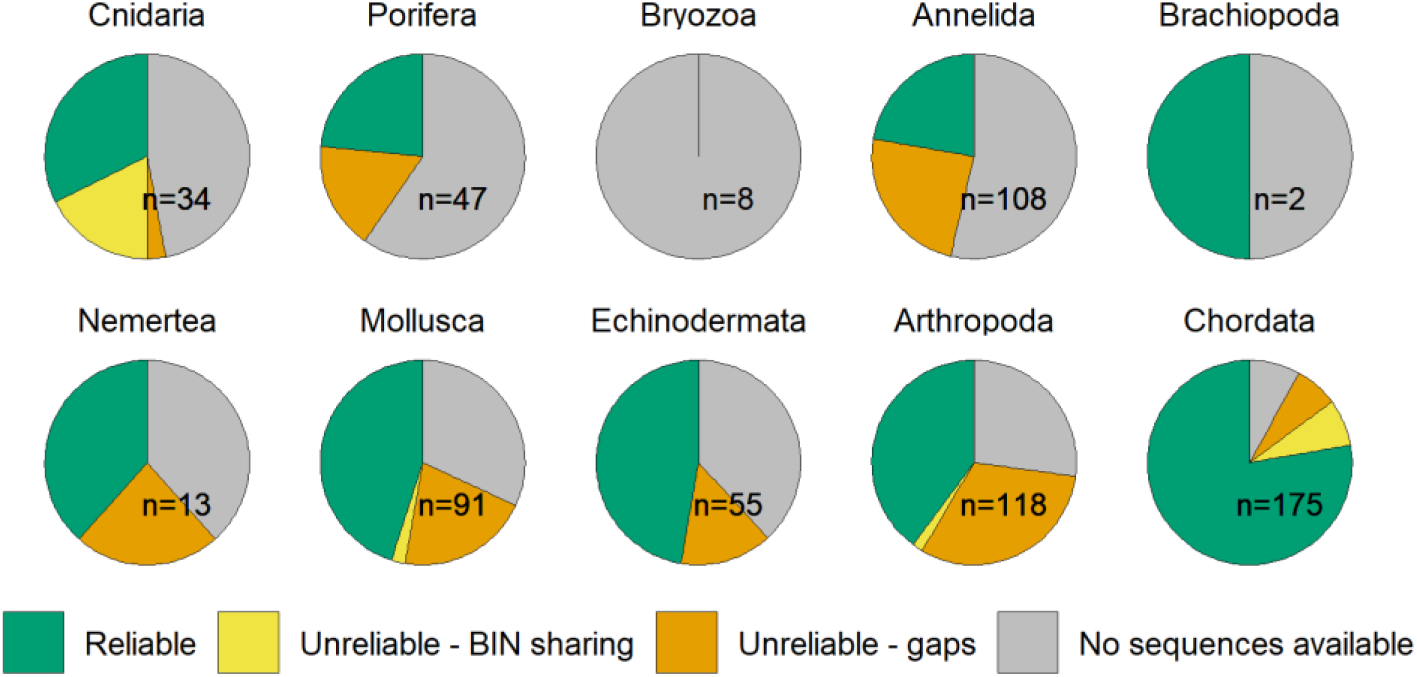
Classification of 651 marine faunal species previously observed in the Gulf of St. Lawrence and included in the GSL-rl, by phylum. Species reliability ranking is based on the availability from local species and sequence similarity to closely related species in the Gulf of St. Lawrence.

We then provided a reliability ranking to each species within the GSL-rl based on the completeness of sequences available (Fig 2, Table S4; see methods for more details). Species within the “Reliable” category accounted for the largest proportion of the species with sequences from the regional library (302 species or 68.8%; 133 vertebrates, 169 invertebrates). Species classified to the “Unreliable due to BIN sharing” and the “Unreliable due to gaps” categories represented 5.2% (23 species; 13 Chordata, 10 invertebrates) and 26.0% (114 species; 12 vertebrates, 102 invertebrates) of the GSL-rl species, respectively. The GSL-rl (version 1.0 and future versions) is available on GitHub (https://github.com/GenomicsMLI-DFO/MLI_GSL-rl).

### Accuracy of species assignments using NCBI-nt and two assignment methods

The proportions of species assignments overall taxa were higher with the Blast-TopHit method (range: 85.5% – 87.9%) than the Blast-LCA method (range: 47.6 – 71.0%) with any identity thresholds (Fig 3A). Overall taxa, the proportions of species assignments increased for the Blast-LCA method while they decreased with the Blast-TopHit method with increasing identity thresholds (Fig. 3A). Across taxonomic groups, the proportions of species assignments were also consistently greater at all thresholds for the Blast-TopHit method compared to the Blast-LCA method. The proportions of species assignments at the 97% similarity threshold varied with the Blast-TopHit from 74.4% for Annelida, Brachiopoda, Nemertea to 93.1% for Arthropoda method and with the Blast-LCA method from 35.8% for Cnidaria, Porifera to 75.2% for Arthropoda (Fig. 3B).

**Figure 3.**
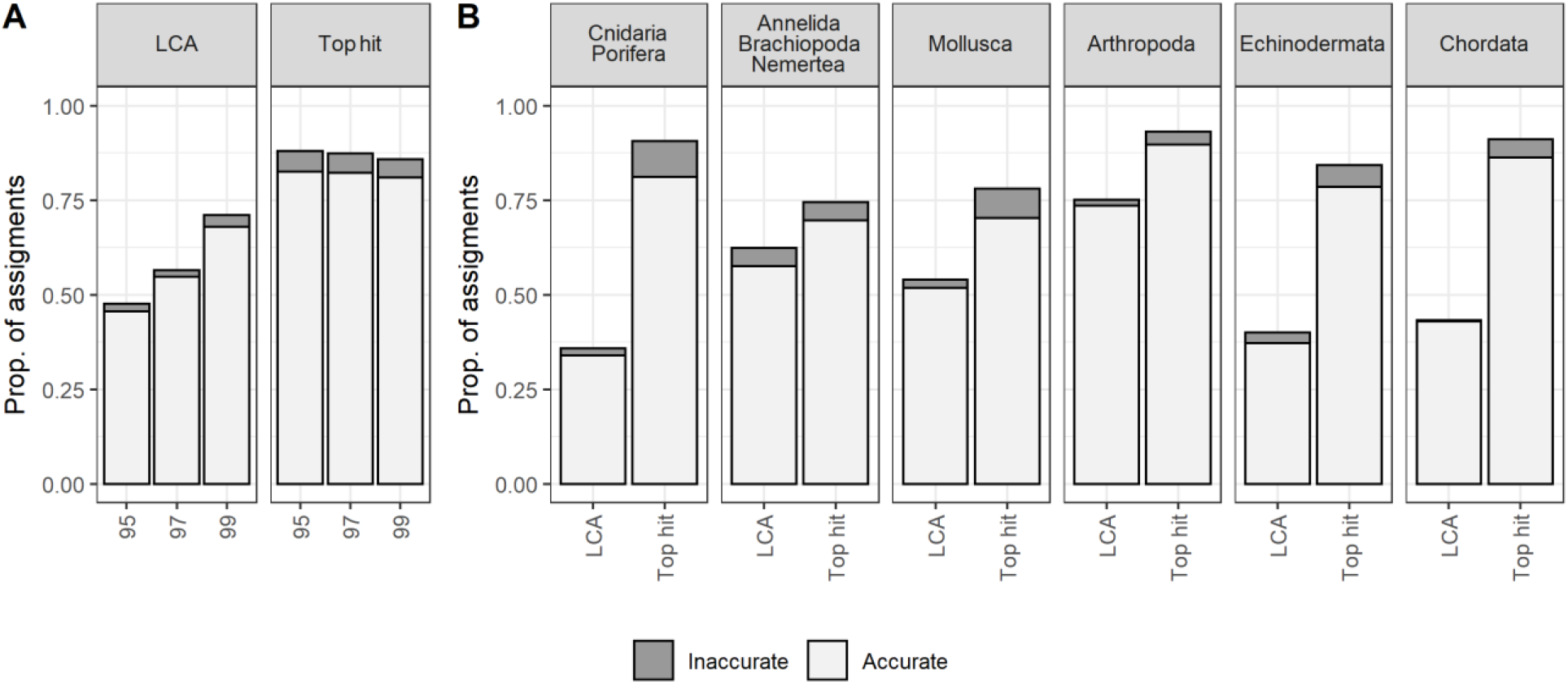
Results of taxonomic assignment of sequences from the GSL-rl using NBCI-nt and Blast-LCA or Blast-TopHit methods. Panels A and B present the proportions of accurate and inaccurate species assignments. Results are presented for all taxonomic groups at the three identity thresholds (95%, 97%, 99%; panel A) and by taxonomic group at the 97% threshold (panel B).

The proportions of accurate species assignments were higher with the Blast-TopHit method compared to the Blast-LCA method, overall taxa and in each taxonomic group at all identify thresholds (Fig. 3A, B). Overall taxa, proportions of accurate species assignments varied between 80.1 and 82.5% for Blast-TopHit method and between 42.7 and 68.0% for the Blast-LCA method for the three identity thresholds tested (Fig. 3A). For each taxonomic group, proportions of accurate assignments at the species level were consistently higher at all thresholds with the Blast-TopHit method compared to the Blast-LCA method. The proportions of accurate species assignments at the 97% threshold varied with the Blast-TopHit method from 69.6% for Annelida, Brachiopoda, Nemertea to 89.6% for Arthropoda and with the Blast-LCA method from 34.0% for Cnidaria and Porifera to 73.6% for Arthropoda (Fig. 3B).

The accuracy was greater for the Blast-LCA method compared to those of the Blast-TopHit method overall taxa at all thresholds (Blast-LCA range: 95.7 – 96.9 %, Blast-TopHit range: 93.8 – 94.4%; Fig. 3A), and in most taxonomic groups at the 97% threshold (Blast-LCA range: 92.3 – 99.2%, Blast-TopHit range: 89.6 – 96.3%; Fig. 3B).

### Species assignments using GSL-rl and NCBI-nt, three assignment methods, and a metabarcoding dataset

We used an eDNA metabarcoding dataset to compare the number and the reliability of species assigned using GSL-rl and NCBI-nt, and three assignment methods. The five possible combinations of repository/library and assignment methods were GSL-rl and NCBI-nt with Blast-LCA (1, 2), GSL-rl and NCBI-nt with Blast-TopHit (3,4), and GSL-rl with IDtaxa (5; Fig 1). A total of 80 species were assigned with the five combinations of repository/library and assignment methods (Fig. 4A). Detected species differed using NCBI-nt and GSL-rl and the three assignment methods (Fig. 4A).

**Figure 4.**
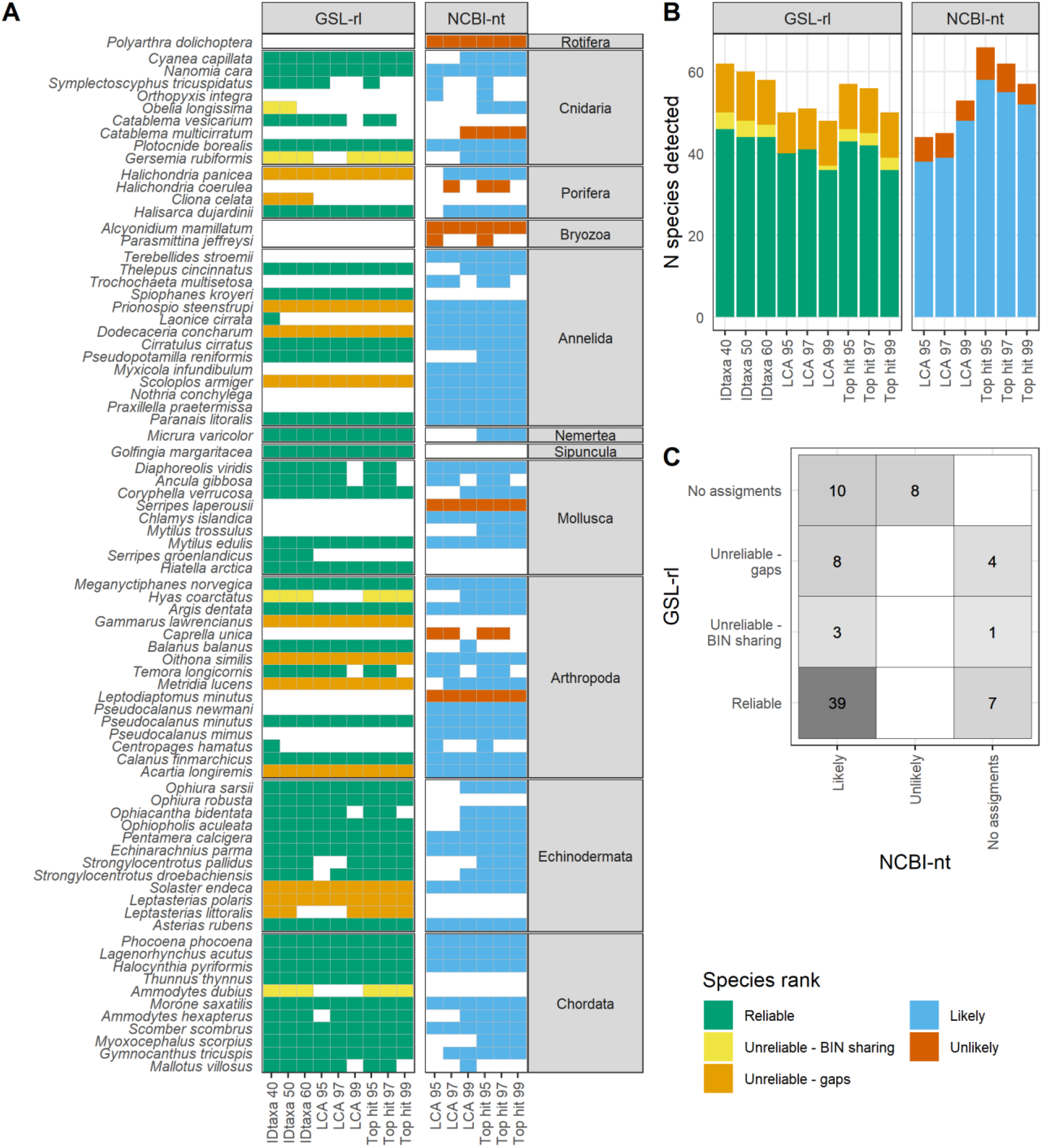
Assignment results at the species level using a regional library (GSL-rl) or a public repository (NBCI-nt), and popular assignment methods. We used three assignment methods, namely IDtaxa (confidence levels: 40%, 50% and 60%), Blast-LCA, and Blast-TopHit (identity thresholds: 95%, 97% and 99%). Panel A detailed the detections for each species, and panel B synthesize the number of species assignments for each source of reference sequences and method. Panel C compared the species rank for all the species assigned with the two sources. Species rank categories are based on sequence availability and sequence similarity to closely related species in the Gulf of St. Lawrence for GSL-rl and on the geographic plausibility for NCBI-nt.

Across all combinations, the highest and lowest numbers of species assigned were observed with NCBI-nt and Blast-TopHit95 (66 species) and Blast-LCA95 (44 species), respectively (Fig. 4B). The number of assigned species decreased with increasing thresholds for most combinations, except for Blast-LCA with the NCBI-nt (Fig. 4B). For GSL-rl, proportions of assigned species ranked as “Unreliable due to BIN sharing” or “Unreliable due to gaps” did not increase or decrease linearly with changing thresholds of assignment methods (Fig. 4B). For NCBI-nt, decreasing proportions of “Unlikely” species were assigned with increasing identity thresholds of Blast-LCA or Blast-TopHit (Fig. 4B).

The assignment method with the maximum number of assigned species differed between GSL-rl and NCBI-nt. The maximum number of assigned species was 62 species with the GSL-rl and IDtaxa40 and 66 species with NCBI-nt and TopHit95 (Fig. 4B). Out of the 62 species assigned using the GSL-rl/IDtaxa40 combination, 46 species (74.2%) were ranked as “Reliable”. The remaining assigned species were ranked as “Unreliable due to BIN sharing” (4 species, 6.5%) or “Unreliable due to gaps” (12 species, 19.4%; Fig. 4B). With the NCBI-nt/TopHit95 combination, 58 (87.9%) and 8 (12.1%) assigned species were ranked as “Likely” and “Unlikely” present, respectively (Fig. 4B).

Large proportions of detected species were exclusively assigned using GSL-rl or NCBI-bt. A total of 30 species (37.5% of all species detected) were assigned only using GSL-rl (12 species) or NCBI-nt (18 species; Fig. 4AC). For the species only assigned with GSL-rl, 7 species were ranked as “Reliable” whereas 1 and 4 species were ranked as “Unreliable due to BIN sharing” and “Unreliable due to gaps”, respectively (Fig. 4AC). For the species only assigned with NCBI-nt, 10 species were considered likely to be present in the GSL whereas 8 species were considered unlikely to be present. For the other 50 species assigned with both GSL-rl and NCBI-nt, 39 species were ranked as “Reliable” with the GSL-rl (78.0%, Fig. 2C). The remaining species assigned belonged to the “Unreliable due to BIN sharing” (3 species, 6.0%) or the “Unreliable due to gaps” categories (8 species, 16.0%; Fig. 4AC).

## Discussion

Species assignments using DNA barcoding and metabarcoding are affected by the quality and taxonomic coverage of reference sequences and assignment methods used. In this study, we first created a regional library (GSL-rl) for a widely used barcoding gene, COI, that includes a reliability ranking system for species level assignments. We then estimated the accuracy of species assignments with NCBI-nt and two assignment methods using the curated sequences from GSL-rl. While Blast-LCA was the most accurate method when using NCBI-nt, the proportion of accurate species assignments was highest with Blast-TopHit (>80 % overall, between 70 and 90% amongst taxonomic groups). We also compared the number and reliability of species assignments using GSL-rl or NCBI-nt and popular assignment methods with a metabarcoding dataset. The reliability of species assignments was the greatest using the regional library but the regional library and the public repository both provided exclusive plausible species detections, highlighting the importance to use both resources as reference databases. Consequently, we also discuss a two-step approach, using first a regional library followed by a public library, to maximize the reliability of species assignments and the number of species detected in metabarcoding studies.

### A COI regional library with a reliability ranking system for metazoans from the Gulf of St. Lawrence (GSL-rl)

The GSL-rl, a curated regional library, provides explicit reliability ranking for 651 species observed within the Gulf of St. Lawrence. We used two simple broad categories, namely “Reliable” and “Unreliable”, to characterize the robustness of species assignments in eDNA metabarcoding studies. This simple ranking system should limit inaccuracies in the interpretation of species assignments for anyone, even with limited scientific background. Past studies have shown the importance of a ranking system to limit erroneous species assignments (e.g., Costa et al. 2012, Knebelsberger et al. 2014). However, the ranking systems used in these studies are targeting an audience of barcoding specialists. With the ranking system of species assignments in GSL-rl, we aimed to keep this classification simple to reach the large audience of eDNA users. Still, we used two “Unreliable” subcategories to highlight 1) the taxa necessitating future barcoding efforts, and 2) the relevancy of the COI barcode to discriminate species. This allows any eDNA scientist to discard the COI loci if species of interest are not discriminated.

The reliability ranking of species in GSL-rl may change over time, particularly for understudied species. The reliability ranking is based on recent information of regional species distributions and intra- and inter-genetic diversity. In the future, species may be upgraded to the “Reliable” category when further sequencing results to fill in data gaps. Some species may also be downgraded to the “Unreliable” category, particularly for complex taxonomic groups in the region that should be targeted for review (e.g., polychaete worms). For these reasons, continuous review, curation, and versioning of any regional libraries are important to track changes in reliability ranking through times. A biennial review is planned for the GSL-rl given the large amount of effort required for such work.

The GSL-rl covers vertebrates and invertebrate species of interest for conservation in the Gulf of St. Lawrence. The GSL-rl contains reference sequences for 439 species of the 651 targeted species in this study (i.e., 67.4%), with reference sequences available for a relatively large proportion of invertebrates (i.e., 59.1%). In Europe, marine invertebrates represented the taxonomic group with the lowest barcode coverage, where only 22.1% had one or more sequences (Weigand et al. 2019). The larger proportion of invertebrates with reference sequences in the GSL-rl is likely due to the species selection to initiate this regional library but also to the smaller study area and the barcoding campaigns for invertebrates in the Northwest Atlantic (e.g., Radulovici et al. 2009, Layton et al. 2016).

The “Reliable” category represented the vast majority of species with reference sequences (68.8%, 302 species) in GSL-rl. Similar results were obtained for marine fish species from Portugal at the COI locus (73.5%, grade A, Costa et al. 2012). About a quarter of the GSL-rl species was ranked as “Unreliable due to gaps” due to missing reference sequences from a relative species. For example, the common shelf species of purple sunstar *Solaster endeca* was only ranked as “Unreliable due to gaps” because no reference sequence was available for the local but rare, deep water congener species, *Solaster earlli*. Generally, taxonomic groups for which fewer sequences were recovered in the GSL-rl were also those with more recognized gaps (e.g., Mollusca, Arthropoda). Furthermore, the GSL-rl is in its early development (v.1.0) and presently covers only a quarter of the estimated 2200 marine faunal species that may occur in the Gulf of St. Lawrence (Nozères 2017). Some groups were more underrepresented than others in GSL-rl, e.g., only 10% of the 250 described amphipods. Our study summarizes and lists where future barcoding efforts should be done to fill gaps in the Gulf of St. Lawrence and the Northwest Atlantic.

The GSL-rl could also improve species assignments in eDNA metabarcoding studies of the Northwest Atlantic and the Arctic Oceans compared to large public databases. The Gulf of St. Lawrence is a transitional marine region where temperate southern species may occur alongside boreal and arctic species (Bourdages et al. 2022). There are no regional libraries covering marine metazoan species at the COI locus in nearby regions, and the GSL-rl could seed the creation of these regional libraries. These can be created by data mining and curating existing sequences from public repositories (e.g., the approach used in this study), completely de novo from barcoding local specimens (e.g., Delrieu-Trottin et al. 2019), or from a combination of both approaches (e.g., Stoeckle et al. 2020, Gold et al. 2021). New tools are now emerging to facilitate the creation of regional reference libraries (e.g., Meta-Fish-lib, Collins et al. 2021; Barcode, Audit & Grade System (BAGS), Fontes et al. 2021). Some of the tools, such as BAGS, even allow for the annotation of species based on concordance between morphological species-based identification and sequence clusters in BOLD (Fontes et al. 2021). Combined with tools to find gaps in reference sequence libraries (e.g., GAPeDNA, Marques et al. 2021), more comprehensive species-level assignments are now possible.

### Accuracy of species assignments using NCBI-nt and two assignment methods

We estimated three performance parameters for metazoan species assignment using NCBI-nt. We observed large variation in those parameters with the two assignment methods tested. While the Blast-LCA method provided overall higher accuracy in species assignments, the proportion of accurate species assignments was greater with Blast-TopHit due to the large difference in the proportion of species assigned, accurate or not. This might be explained by the sensitivity of both methods to the prevalence of BIN sharing, gaps, and mislabeling within public repositories. For instance, Blast-LCA method is expected to be less precise and cause under-classification (i.e., assignments at a higher taxonomic level) in the presence of closely related species and BIN sharing, lowering the proportion of assignments at the species level. In contrast, Blast-TopHit is expected to favor assignments at the species level and is more impacted by gaps and mislabeled sequences (e.g., Schenekar et al. 2020), lowering the accuracy of species assignments. The relatively good performance of Blast-TopHit observed here suggests that gaps and misidentified specimens within NCBI-nt are limited for the targeted marine species of the Gulf of St. Lawrence.

Previous studies have shown that assignment methods can affect taxa detected in metabarcoding studies (O’Rourke et al. 2020, Hleap et al. 2021). Our results confirmed those from Hleap and colleagues (2021) that the Blast-TopHit method outperformed the Blast-LCA method with NCBI-nt to provide higher proportion of assignments. Our results also show that the proportion of accurate species assignments varied largely between taxonomic groups. Relatively well-described marine taxonomic groups such as Arthropoda (i.e., crustaceans) and Chordata (i.e., fishes and mammals) have reached proportions of accurate species assignments ≥85% with the Blast-TopHit method. The proportion of accurate species assignments is much lower for the Blast-LCA approach with the Chordata (43%), probably because this approach is more sensitive to the presence of close relative species (i.e., BIN sharing) reducing the potential of species level identification. Other groups such as Annelida, Brachiopoda, Nemertea, and Mollusca achieved lower proportions of accurate species assignments using both Blast-TopHit and Blast-LCA methods (maximum 70%).

The proportion of accurate species assignments using the sequences from the GSL-rl in our study will be different at the time of reading this article due to the continuous growth of the public repository NCBI-nt. The publication of new sequences of low quality or with incorrect species identification can create unexpected ambiguities in species assignments as public repositories grow (Locatelli et al. 2020, Radulovici et al. 2021). Without a comprehensive versioning system, changes in the NCBI-nt database also limit the reproducibility of species assignments as it is difficult to identify and to access a specific release. Note that starting with Blast v.2.13 launched in March 2022, it is now possible to generate a metadata file describing the database used (Camacho and Madden 2022), which in an important step toward higher traceability.

### Comparing reliability of species assignments using GSL-rl and NCBI-nt, three assignment methods, and a metabarcoding dataset

The method with the maximal number of species assigned in the metabarcoding dataset differed between GSL-rl and NCBI-nt. The IDtaxa40 assignment method provided the highest number of species assigned using the GSL-rl. Sequence composition strategies for species assignments such as IDtaxa, QIIME, and RDP, had contrasting performance results in previous benchmarking studies (O’Rourke et al. 2020, Hleap et al. 2021, Mathon et al. 2021). Our results contrast with those from a previous study showing that IDtaxa did not perform as well as Blast with mock communities composed of various freshwater taxonomic groups (Hleap et al. 2021). The contrasting results between the latter and our studies could be explained by the difference in the confidence threshold used (Hleap et al. 2021). Parameter tuning while using any approach may be key to choosing an optimal method for a dataset while more benchmarking studies are undertaken. The relatively better performance of IDtaxa in our study might also be due to the quality of the regional library used to train the classifier. We know little about the impact of using training sets of different qualities on taxonomic assignment using classifiers, and the gains of using regional libraries might be important in this context. With NCBI-nt, the number of detected species was greater with Blast-TopHit compared to Blast-LCA with the metabarcoding dataset. Those results are similar as those obtained with the GSL-rl COI sequences and have been discussed in the previous section.

More than a third of the species assigned (n = 33 out of 80) were exclusive to the use of GSL-rl or NCBI-nt with the metabarcoding dataset. For the GSL-rl, the exclusion of non-indigenous species or mislabeled sequences increased the number of species assigned, confirming previous studies results improving species assignments with regional libraries (von Ammon et al. 2018, Gold et al. 2021). The exclusion of non-indigenous species increased the taxonomic resolution with the GSL-rl of the Atlantic bluefin tuna *Thunnus thynnus*. Underclassification is usually observed when using NCBI-nt as the Atlantic bluefin tuna presently shares a BIN (BOLD: AAA7352) with other *Thunnus* species that are not expected to be present in the Gulf of St. Lawrence (Nozères 2017). With NCBI-nt, detections in the “Likely” category comprised species for which sequences were not included in the GSL-rl because of the stringency of quality filtering performed (e.g., Iceland scallop *Chlamys islandica*). Other detections in the “Likely” category were included in the GSL-rl (e.g., the polychaete worm *Terebellides stroemii*) but the inability to detect them suggests that their intra-specific diversity is not fully covered by the GSL-rl. Finally, a few species assigned with NCBI-nt were not listed as present in the GSL but after reconsideration are likely to be found in the target area (e.g., *Pseudocalanus newmani*). With NCBI-nt, we also observed underclassification of the sea star genus *Leptasterias* due to sequence mislabeling, which is expected to occur (Bidartondo 2008, Mioduchowska et al. 2018). The underclassification is due to two misidentified sequences, one is for *Leptasterias littoralis* identified as the sea star *Asterias forbesi* and the other is for *Leptasterias polaris* identified as the butterfly *Polyommatus fulgens*.

Contrasting the ranking category of NCBI-nt and GSL-rl revealed an important gain in reliability with our annotated regional library (Fig 1). With NCBI-nt, we provided the likeliness of a species to be present in the Gulf of St. Lawrence due to the availability of a public species list (Nozères 2017). Such information layer is often difficult to obtain without expert knowledge (Pappalardo et al. 2021). Of all the species ranked in the “Likely” category using NCBI-nt, around 78% were classified as “Reliable” in the GSL-rl. Our results showed that the remaining 22% should be interpreted with caution given gaps (16%) or BIN sharing with close relative species (6%; Fig 4C). Our results suggest that species level assignments of a metabarcoding dataset using NCBI-nt and a filter based on geographic plausibility can be misleading. This important hidden and overlooked uncertainty could be acceptable for empirical studies but not under regulatory context where specific species identification can be crucial, such as the identification of species at risk (Gilbey et al. 2021). Evaluation of false-positives in the detections of endangered or invasive species should include potential bias caused by gaps in reference libraries (Cristescu and Hebert 2018).

### Maximizing the reliability and the number of species assignments in eDNA metabarcoding studies using a regional library and a public repository

Our results showed that the use of a regional library increase both the reliability and the number of species detected with an eDNA metabarcoding dataset. Yet, some species likely present in the Gulf of St. Lawrence were only detected with NCBI-nt, as discussed in the previous section. This will be a recurrent problem until reference sequences are available for more marine metazoan species in the Gulf of St. Lawrence. The growth of GSL-rl will increase the number of species that can be detected using the regional library but unexpected species, such as new invasive species or species that have recently expanded their distribution, could remain undetected (Bohmann et al. 2014, Klymus et al. 2017, Piper et al. 2019, Stoeckle et al. 2020, Gold et al. 2021). Restricting species assignments to GSL-rl and avoiding the use of NCBI-nt would limit the maximum number of species detected.

Combining the strengths of a regional library and public repositories as a two-step approach is consequently the optimal solution to maximize reliability and the number of species assigned in metabarcoding studies. Taxonomic assignments should be first performed with a regional library, ideally including a reliability ranking system as in the GSL-rl, to maximize the confidence in species assignment. We then strongly advise contrasting species assignment results from a regional library with those using a public repository to increase the number of species detections (see also Piper et al. 2019, and Xiong et al. 2022 for similar recommendations). This would allow the reader to have a qualitative estimation of the proportion of accurate species assignments. Species assignments relying uniquely on NCBI-nt should also clearly indicate that their reliability is limited.

We also encourage further benchmarking studies for the selection of optimal methods based on a broader comparison of assignment methods and the development of training sets for machine-learning methods. We limited the size of this study by selecting assignment methods often used in eDNA metabarcoding studies that are also performing relatively well in benchmarking assignment studies (O’Rourke et al. 2020, Hleap et al. 2021). Our results emphasize that future benchmarking studies should be done independently for regional libraries and public repositories, given the different properties of these resources, and to maximize the reliability and the number of species assignments.

## Acknowledgements

We thank Grégoire Cortial and Jade Larivière for their inputs at the earlier stages of this study. We also thank Nick Jeffery for helpful comments on a previous version of the manuscript. We thank Yanick Gendreau and Sandra Velasquez from the Coastal environmental baseline program and Geneviève Faille and Geneviève Côté from the Banc-des-Américains Marine Protected Area for eDNA sampling and the initial list of marine faunal species of interest.

## Data Accessibility

The data and scripts used in this manuscript are stored in the github repository https://github.com/GenomicsMLI-DFO/GSL_COI_ref_library

The GSL-rl (sequences, reliability ranking and trained dataset) can be found in the github repository https://github.com/GenomicsMLI-DFO/MLI_GSL-rl

## Benefit-Sharing

Benefits from this research accrue from the sharing of our data and results on public databases as described above.

## Author Contributions

A.B. co-conceived and co-developed the ideas underlying the manuscript, wrote scripts, compiled GSL-rl, analyzed data, and co-wrote the manuscript. C.N. initiated the project of a reference library for the GSL, co-conceived and co-developed the ideas, revised GSL-rl, analyzed data and edited all drafts. E.P. initiated the project of a reference library for the GSL and revised the manuscript. G.J.P. co-conceived and co-developed the ideas, co-wrote the manuscript and secured funding.

