## supplementary material for "Maximizing the reliability and the number of species assignments in metabarcoding studies"

### A. Supplementary figures and tables from the manuscript

Fig. S1 – Steps to obtain, filter and select publicly available sequences
Table S1 – Metadata of the GSL-rl, including ranking systems. [csv file]
Table S2 – BINs shared by two or more taxa
Table S3 – Taxa sharing more than one BIN within the GSL-rl
Table S4 – Characteristic of the curated regional library of the Gulf of St. Lawrence (GSL-rl) version 1.0.

### B. Creation of Gulf of St. Lawrence regional library (GSL-rl)

### C. Creation of an eDNA metabarcoding dataset

### A. Supplementary figures and tables from the manuscript

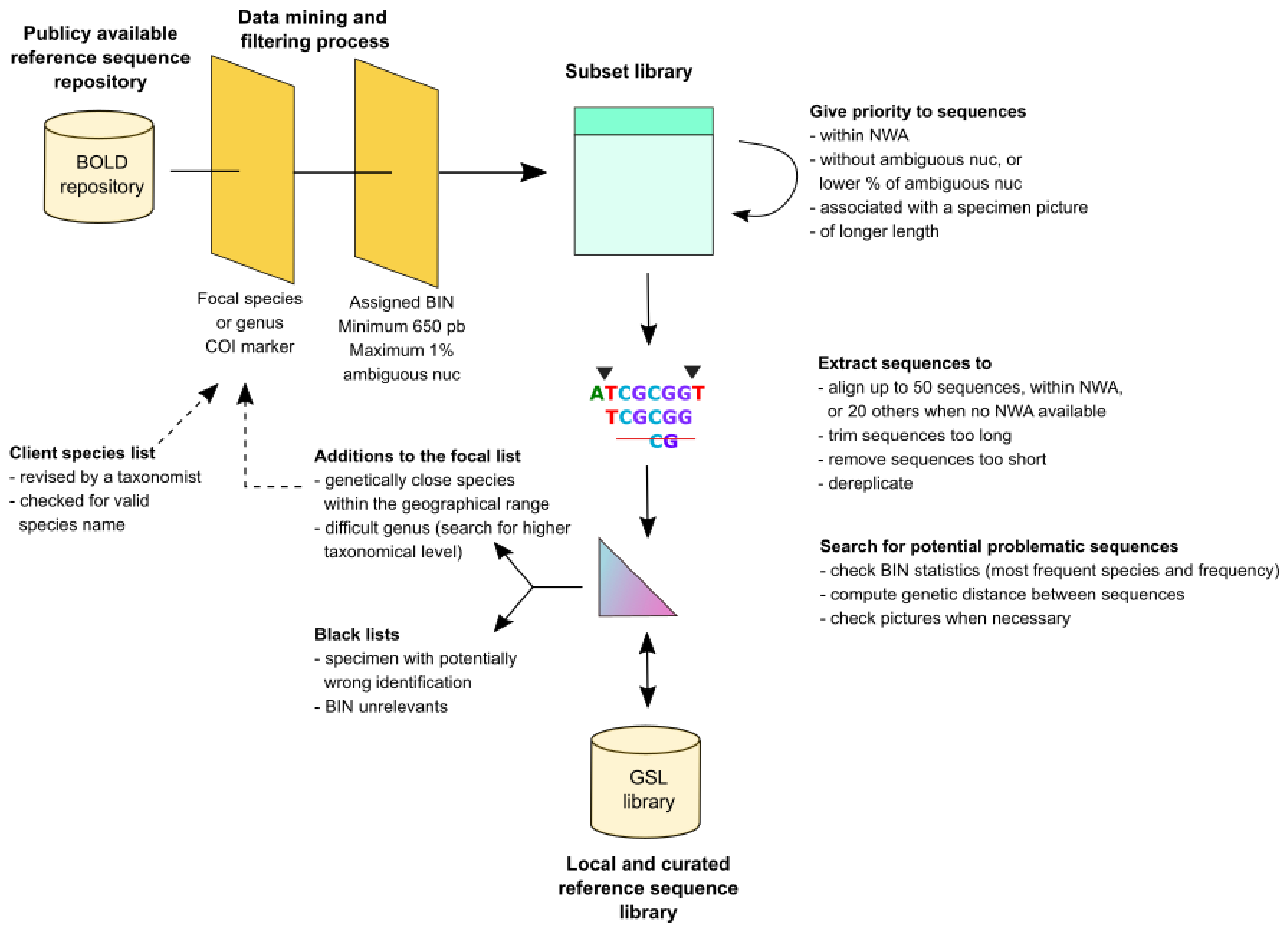


**Figure S1 Summary of the steps to obtain, filter and select publicly available sequences to create a regional library.** First, a filtration step through BOLD metadata. Second, remaining sequences are ordered based on various criteria. Then sequences are extracted, aligned, trimmed, filtered and dereplicated. Remaining sequences are checked to identify potential identification problems, from BIN information and genetic distances, then added to the curated regional library for the Gulf of St. Lawrence (GSL-rl). Since it is an iterative process, new species or genus can be searched again to further improve the GSL-rl. All these steps can be done through an R code.

**Table S2.** **BINs shared by two or more taxa in the GSL-rl**

| Category | Phylum | BIN | Species |
|---|---|---|---|
| Vertebrate | Chordata | BOLD:AAB1380 | *Alosa aestivalis*, *Alosa pseudoharengus* |
| Vertebrate | Chordata | BOLD:AAB4332 | *Ammodytes americanus*, *Ammodytes dubius* |
| Vertebrate | Chordata | BOLD:AAA9928 | *Aspidophoroides monopterygius*, *Aspidophoroides olrikii* |
| Vertebrate | Chordata | BOLD:AAB3781 | *Eumicrotremus spinosus* †, *Eumicrotremus terraenovae* |
| Vertebrate | Chordata | BOLD:ACA5464 | *Gymnelus hemifasciatus* ‡, *Gymnelus viridis* |
| Vertebrate | Chordata | BOLD:AAB4897 | *Liparis atlanticus*, *Liparis inquilinus* ‡ |
| Vertebrate | Chordata | BOLD:AAC3262 | *Lycodes lavalaei*, *Lycodes polaris* ‡ |
| Vertebrate | Chordata | BOLD:ACE9999 | *Sebastes fasciatus*, *Sebastes norvegicus*, *Sebastes mentella* |
| Invertebrate | Arthropoda | BOLD:AAA9922 | *Hyas coarctatus* †, *Hyas alutaceus* |
| Invertebrate | Cnidaria | BOLD:AAH8240 | *Clavularia borealis* †, *Alcyonium siderium*, *Gersemia rubiformis* |
| Invertebrate | Cnidaria | BOLD:AAF8957 | *Hormathia nodosa*, *Hormathia digitata* |
| Invertebrate | Cnidaria | BOLD:AAE6029 | *Obelia longissima*, *Obelia dichotoma* |
| Invertebrate | Echinodermata | BOLD:AAB2709 | *Leptasterias littoralis* †, *Leptasterias groenlandica* |
| Invertebrate | Mollusca | BOLD:ACR2733 | *Buccinum hydrophanum* ‡, *Buccinum scalariforme* |
| Invertebrate | Mollusca | BOLD:AAF2554 | *Nuculana tenuisulcata*, *Nuculana pernula* |
| Invertebrate | Porifera | BOLD:ACQ6272 | *Polymastia thielei*, *Polymastia hemisphaerica* |

† : species likely absent from region

‡ : species status uncertain

**Table S3 Taxa sharing more than one BIN within the GSL-rl**

| Category | Phylum | Taxon | N BIN | BIN |
|---|---|---|---|---|
| Vertebrate | Chordata | *Alepocephalus bairdii* | 2 | BOLD:AAD7907, BOLD:ABY4903 |
| Vertebrate | Chordata | *Chauliodus sloani* | 2 | BOLD:AAB1178, BOLD:AAB1179 |
| Vertebrate | Chordata | *Liparis inquilinus* | 2 | BOLD:AAB4897, BOLD:ACA5502 |
| Vertebrate | Chordata | *Lycodes spp* | 2 | BOLD:ABZ0973, BOLD:ACE4278 |
| Vertebrate | Chordata | *Mola mola* | 3 | BOLD:ADK1672, BOLD:AAD0035, BOLD:AAD0034 |
| Vertebrate | Chordata | *Synaphobranchus kaupii* | 2 | BOLD:AAA8286, BOLD:AAA8287 |
| Invertebrate | Annelida | *Aphelochaeta filiformis* | 2 | BOLD:ACH1048, BOLD:ACH1047 |
| Invertebrate | Annelida | *Bradabyssa villosa* | 4 | BOLD:ACH1761, BOLD:ABZ8261, BOLD:AAW7084, BOLD:ACM2001 |
| Invertebrate | Annelida | *Gattyana cirrhosa* | 2 | BOLD:AAC0429, BOLD:AAG5103 |
| Invertebrate | Annelida | *Harmothoe imbricata* | 3 | BOLD:ABZ7769, BOLD:AAB1625, BOLD:AAA9124 |
| Invertebrate | Annelida | *Laonice cirrata* | 3 | BOLD:AAD5360, BOLD:AAU3353, BOLD:AAW9888 |
| Invertebrate | Annelida | *Limnodrilus hoffmeisteri* | 4 | BOLD:ACB6505, BOLD:AAA6803, BOLD:ADX1968, BOLD:ACM1418 |
| Invertebrate | Annelida | *Nicolea zostericola* | 2 | BOLD:AAC0291, BOLD:ADW6705 |
| Invertebrate | Annelida | *Pista cristata* | 2 | BOLD:AAH9500, BOLD:ACP4603 |
| Invertebrate | Annelida | *Polycirrus medusa* | 3 | BOLD:ACE5004, BOLD:AAF1233, BOLD:ABY4561 |
| Invertebrate | Annelida | *Potamothrix moldaviensis* | 2 | BOLD:ACK5400, BOLD:ACK5110 |
| Invertebrate | Annelida | *Terebellides spp* | 2 | BOLD:AAD6167, BOLD:AAF2172 |
| Invertebrate | Annelida | *Terebellides stroemii* | 3 | BOLD:ACP0580, BOLD:AAE9393, BOLD:AAE9392 |
| Invertebrate | Annelida | *Tubifex tubifex* | 7 | BOLD:ADJ9700, BOLD:ADK4845, BOLD:AAA6801, BOLD:ADX0052, BOLD:AAA6802, BOLD:ACV8476, BOLD:ACA9222 |
| Invertebrate | Arthropoda | *Acartia clausi* | 3 | BOLD:ACB5005, BOLD:ACB4079, BOLD:AAT9961 |
| Invertebrate | Arthropoda | *Acartia hudsonica* | 2 | BOLD:AAJ3150, BOLD:ACL9285 |
| Invertebrate | Arthropoda | *Acartia tonsa* | 2 | BOLD:AAA5450, BOLD:AAA5453 |
| Invertebrate | Arthropoda | *Anonyx spp* | 2 | BOLD:AAF0743, BOLD:AAA9156 |
| Invertebrate | Arthropoda | *Anonyx ochoticus* | 2 | BOLD:ACA9971, BOLD:ADF0772 |
| Invertebrate | Arthropoda | *Argis dentata* | 2 | BOLD:ADK3850, BOLD:AAC3186 |
| Invertebrate | Arthropoda | *Calanus finmarchicus* | 2 | BOLD:ACP6217, BOLD:ACP5847 |
| Invertebrate | Arthropoda | *Cancer irroratus* | 2 | BOLD:AAB7738, BOLD:ACD1651 |
| Invertebrate | Arthropoda | *Centropages typicus* | 2 | BOLD:AAW6024, BOLD:ACM7827 |
| Invertebrate | Arthropoda | *Chionoecetes opilio* | 2 | BOLD:AAA3656, BOLD:ABZ6341 |
| Invertebrate | Arthropoda | *Crangon septemspinosa* | 2 | BOLD:AAB5942, BOLD:ACL7976 |
| Invertebrate | Arthropoda | *Eualus gaimardii* | 2 | BOLD:AAE4771, BOLD:AAJ0881 |

| | | | | |
|---|---|---|---|---|
| Invertebrate | Arthropoda | *Gammarus setosus* | 4 | BOLD:ABZ4044, BOLD:ACE3104, BOLD:AAA3651, BOLD:AAA3653 |
| Invertebrate | Arthropoda | *Gammarus tigrinus* | 3 | BOLD:ADP7774, BOLD:ADP7773, BOLD:AAA4302 |
| Invertebrate | Arthropoda | *Ischyrocerus anguipes* | 2 | BOLD:AAC5063, BOLD:AAC0651 |
| Invertebrate | Arthropoda | *Lebbeus polaris* | 6 | BOLD:AAB4526, BOLD:AAG9842, BOLD:AAB4528, BOLD:ACF1644, BOLD:AAB4529, BOLD:AAB4527 |
| Invertebrate | Arthropoda | *Nymphon stroemi* | 2 | BOLD:ADF4183, BOLD:AAG4693 |
| Invertebrate | Arthropoda | *Oithona similis* | 3 | BOLD:AAG5172, BOLD:ACK9027, BOLD:ADU0358 |
| Invertebrate | Arthropoda | *Spirontocaris spinus* | 4 | BOLD:AAB6143, BOLD:AAB6146, BOLD:AAB6144, BOLD:ACR1068 |
| Invertebrate | Cnidaria | *Cyanea capillata* | 2 | BOLD:AAP1190, BOLD:AAD3480 |
| Invertebrate | Echinodermata | *Gorgonocephalus arcticus* | 2 | BOLD:AAC8875, BOLD:ABY6858 |
| Invertebrate | Echinodermata | *Gorgonocephalus eucnemis* | 3 | BOLD:AAI5449, BOLD:AAE7308, BOLD:ACQ0423 |
| Invertebrate | Echinodermata | *Henricia* spp | 4 | BOLD:AAD3463, BOLD:AAD3482, BOLD:AAB3569, BOLD:AAB9183 |
| Invertebrate | Echinodermata | *Leptasterias polaris* | 2 | BOLD:ACE6383, BOLD:ADV1115 |
| Invertebrate | Echinodermata | *Ophioscolex glacialis* | 2 | BOLD:ACV6677, BOLD:ACV6676 |
| Invertebrate | Echinodermata | *Ophiura sarsii* | 2 | BOLD:ACO7183, BOLD:AAD3481 |
| Invertebrate | Echinodermata | *Pseudarchaster parelii* | 2 | BOLD:AAD5295, BOLD:AAH8175 |
| Invertebrate | Mollusca | *Ancula gibbosa* | 2 | BOLD:ADL9788, BOLD:ACW0116 |
| Invertebrate | Mollusca | *Buccinum hydrophanum* | 2 | BOLD:ACR2733, BOLD:ADE0634 |
| Invertebrate | Mollusca | *Cryptonatica affinis* | 7 | BOLD:ADY4343, BOLD:AAO4021, BOLD:ADS1898, BOLD:ADE1312, BOLD:ABA4155, BOLD:ADY4344, BOLD:ADF1526 |
| Invertebrate | Mollusca | *Euspira heros* | 2 | BOLD:ABW0896, BOLD:ABW0528 |
| Invertebrate | Mollusca | *Hiatella arctica* | 2 | BOLD:ABA8494, BOLD:ADC8266 |
| Invertebrate | Mollusca | *Lepeta caeca* | 2 | BOLD:AAX5488, BOLD:ACB8623 |
| Invertebrate | Mollusca | *Mytilus edulis* | 2 | BOLD:AAA2184, BOLD:AAA2185 |
| Invertebrate | Mollusca | *Nuculana minuta* | 3 | BOLD:ADM2144, BOLD:AEF0228, BOLD:ACM0814 |
| Invertebrate | Mollusca | *Serripes groenlandicus* | 2 | BOLD:AAH9555, BOLD:AAH9554 |
| Invertebrate | Nemertea | *Cephalothrix spiralis* | 2 | BOLD:ACQ6349, BOLD:AAG3611 |
| Invertebrate | Porifera | *Cliona celata* | 3 | BOLD:ACH4809, BOLD:AAO8118, BOLD:ACH3517 |
| Invertebrate | Porifera | *Halichondria panicea* | 2 | BOLD:ABV8568, BOLD:ADX0707 |

**Table S4. Characteristics of the curated regional library of the Gulf of St. Lawrence (GSL-rl) version 1.0.** Number (N) of targeted species in the improved decision maker list and number (N) of targeted species with COI sequences for each of the three reliability categories, namely “Reliable”, “Unreliable due to BIN sharing” and “Unreliable due to gaps”.

| **Phylum** | **Class** | **N species** | **N species with COI sequences** | | | |
|---|---|---|---|---|---|---|
| | | | Reliable | Unreliable due to BIN sharing | Unreliable due to gaps | Total |
| **Cnidaria** | Anthozoa | 24 | 4 | 4 | 1 | 9 |
| | Hydrozoa | 8 | 5 | 2 | 0 | 7 |
| | Scyphozoa | 1 | 1 | 0 | 0 | 1 |
| | Staurozoa | 1 | 1 | 0 | 0 | 1 |
| **Porifera** | Calcarea | 7 | 0 | 0 | 0 | 0 |
| | Demospongiae | 39 | 11 | 0 | 8 | 19 |
| | Hexactinellida | 1 | 0 | 0 | 0 | 0 |
| **Bryozoa** | Gymnolaemata | 8 | 0 | 0 | 0 | 0 |
| **Annelida** | Clitellata | 8 | 4 | 0 | 2 | 6 |
| | Polychaeta | 99 | 19 | 0 | 24 | 43 |
| **Brachiopoda** | Rhynchonellata | 2 | 1 | 0 | 0 | 1 |
| **Nemertea** | Hoplonemertea | 4 | 3 | 0 | 0 | 3 |
| | Palaeonemertea | 1 | 1 | 0 | 0 | 1 |
| | Pilidiophora | 8 | 1 | 0 | 3 | 4 |
| **Sipuncula** | Sipunculidea | 1 | 1 | 0 | 0 | 1 |
| **Mollusca** | Bivalvia | 24 | 9 | 2 | 4 | 15 |
| | Gastropoda | 62 | 30 | 0 | 14 | 44 |
| | Polyplacophora | 5 | 2 | 0 | 1 | 3 |
| **Arthropoda** | Hexanauplia | 22 | 12 | 0 | 7 | 19 |
| | Malacostraca | 81 | 34 | 2 | 29 | 65 |
| | Pycnogonida | 15 | 1 | 0 | 1 | 2 |
| **Echinodermata** | Asteroidea | 31 | 12 | 0 | 6 | 18 |
| | Echinoidea | 3 | 3 | 0 | 0 | 3 |
| | Holothuroidea | 3 | 3 | 0 | 0 | 3 |
| | Ophiuroidea | 18 | 8 | 0 | 2 | 10 |
| **Chordata** | Actinopterygii | 150 | 116 | 13 | 12 | 141 |
| | Appendicularia† | 4 | 0 | 0 | 0 | 0 |
| | Ascidiacea† | 4 | 3 | 0 | 0 | 3 |
| | Elasmobranchii | 12 | 12 | 0 | 0 | 12 |
| | Mammalia | 3 | 3 | 0 | 0 | 3 |
| | Myxini | 1 | 1 | 0 | 0 | 1 |
| | Petromyzonti | 1 | 1 | 0 | 0 | 1 |

†Invertebrates included within the phylum Chordata

### B. Creation of Gulf of St. Lawrence regional library (GSL-rl)

The creation of regional library for the Gulf of St. Lawrence (GSL) metazoans was composed of three major steps: 1) obtaining a preliminary list of species from decision-makers, management personnel and regional taxonomic information, 2) creating an initial regional library using bioinformatics tools and round of reviews (hereafter draft regional library), and 3) enhancing the draft regional library using a metabarcoding dataset (hereafter GSL regional library (GSL-rl); Fig. S1).

#### *Step 1. Reviewed species list*

Decision-makers and management personnel provided a list of taxa (species or higher taxonomic ranks), observed with traditional survey methods, or suspected being present. We enhanced this list by adding potential species within this group present in the geographical range based on a checklist of marine animal species of the GSL (Nozères 2017). This list was reviewed by Claude Nozères, a regional marine biodiversity expert, to validate their presence within the GSL and to highlight species with identification problems at the morphological level or in need of further taxonomic studies. The species names list was then matched with the World Register of Marine Species (WoRMS, WoRMS Editorial Board 2020) through an in-house R script. Recent name changes were noted to allow sequence searches with invalid retrieved names (Table S5). Taxonomic classification hierarchy for each focal taxon was also retrieved using WoRMS database.

#### *Step 2. Draft regional library*

We then used BOLD to seek reference sequences due to the availability of data for COI, the quality of metadata, and the relative ease to access and obtain the data with bioinformatics tools. In BOLD, large metadata are available with each sequence (e.g., regions, sites, pictures of the specimen, sequencing traces), and a BIN information (i.e., unique identifier of sequences based on genetic distance) is also provided, which are not available while working with NCBI-nt.

We built an R script using the package bold (v.1.1.0, Chamberlain 2020) for datamining of COI sequences and metadata for species of interest within BOLD. Searches on BOLD were done with the function *bold_seqspec*. We filtered sequences based on different criteria including markercode as "COI-5P", less than 1% ambiguous sites over the nucleotides, a preference for sampling site between 35–78 latitude and -78–-40 longitude, province_state within Quebec, Nova Scotia, New Brunswick, Newfoundland and Labrador, Prince Edward Island, Maine, New Hampshire, Massachusetts, Rhode Island, Connecticut, New Jersey, Delaware, Maryland, Virginia, column "region", "sector" and "exactsite" including St-Laurent, Saint-Laurent, St Lawrence, St. Lawrence, Fundy, Flemish Cap, Maritimes, Scotian Shelf, Georges Bank, Grand Bank, Baffin Bay, Durban Harbour, but excluding Hudson Bay, Hudson Strait, Beaufort, Nord-du-Quebec, Ungava. Entries without bin_uri defined were excluded. Sequences were classified based on their geographic origin (i.e., NWA or not) and a proxy of quality (i.e., proportion of ambiguous nucleotides), the presence of a picture, and the length of the sequences. When Northwest Atlantic sequences were available, up to 50 were kept to largely cover the genetic diversity. When none were within the geographical limits, up to 20 were kept from undefined or other locations. Sequences were then aligned and marker sites (LCO1490, HCO2198) were retrieved and trimmed before dereplication. Sequences shorter than 650 pb were removed. BIN information for each sequences were kept. For each BIN, a reverse search was made to check all associated taxonomic classifications; whenever discrepancy was observed with taxa-BIN identification (e.g., most sequences associated with another species), further investigations were made to assess the causes (i.e., potential error or genetic similarity to another species; Table S6).

We used the list of species to retrieve the vast majority of sequences. For species with no matching sequences but known recent name change, a search was made with the previous known name (Table S5). Sequences from other species than those from the list were also added if we identified a close relative species present in an area close to the GSL. Finally, for flagged species (i.e., identified a priori as potentially problematic, see Step 1), searches were also made at the genus levels, to retrieve potential sequences within NWA not previously covered by a BIN. We also assessed the number of intraspecific BIN and the number of species within BIN. We computed a genetic distance intra and interspecific between BINs as the Kimura's 2-parameters distance (Kimura 1980) with ape R package (v.5.0, Paradis and Schliep 2019).

*Step 3. GSL regional library*

To check for missing important species from the draft regional library, ESV observed within the metabarcoding dataset (see section C below on the creation of the eDNA metabarcoding dataset) were compared over NCBI-nt (downloaded 2020-10-23), using the BlastX tool *blastn* (v.2.10.1. Camacho et al. 2009). Sequences with at least 95% identity, and with an alignment length>= 95% of the compared sequence, were kept. We examined sequence names within the Metazoa kingdom to detect potential uncovered taxa (i.e., taxa not in the a priori taxa list but possibly within the NWA). New taxa were used for a second round of data mining on BOLD, as previously described in the draft-local library section above. Resulting aligned, trimmed, and filtered sequences were joined to the draft regional library to create an improved GSL regional library (GSL-rl) based on metabarcoding data results.

Each species within the GSL-rl was annotated based on the completeness and similarity to close relative species. Species sharing a BIN group with other species within the GSL-rl were annotated as "Unreliable due to BIN sharing", species close to another species (i.e., within the same genus) with no reference sequences were classified as "Unreliable due to gaps"; all other species, with unique BIN and reference sequence for all known local species from the same genus, were noted "Reliable" (Table S1).
Searches on BOLD and WORMS for the step 2 were performed between 2020-09-30 and 2020-10-05, and between 2021-01-16 and 2021-01-20 for the step 3. Note that *Aulactina stella* was also resolved by hand, because there was an error in the loading of BOLD metadata.

**Table S5 List of species retrieved in BOLD under different names following WoRMS current taxonomy (consulted between 2020-09 and 2021-01).**

| Retrieved name (BOLD) | Current name (WoRMS 2021) | **Reason** |
|---|---|---|
| *Myxine glutinosa* | *Myxine limosa* | Other is NE Atlantic only |
| *Flabellina salmonacea* | *Ziminella salmonacea* | Synonym |
| *Melita dentata* | *Megamoera dentata* | Synonym |
| *Dendronotus niveus* | *Dendronotus elegans* | Synonym |
| *Palaemontes vulgaris* | *Palaemon vulgaris* | Synonym |
| *Palaemonetes pugio* | *Palaemon pugio* | Synonym |
| *Radiella hemisphaerica* | *Polymastia hemisphaerica* | Synonym |
| *Anonyx lilljeborgii* | *Anonyx lilljeborgi* | Synonym |
| *Brada villosa* | *Bradabyssa villosa* | Synonym |
| *Diplocirrus longisetosus* | *Saphobranchia longisetosa* | Synonym |
| *Scoloplos acutus* | *Leitoscoloplos acutus* | Synonym |
| *Halisarca dujardini* | *Halisarca dujardinii* | Synonym |

**Table S6 List of taxa BIN number removed for a specific species from the regional library during the two rounds of data mining on BOLD, and the reason of their exclusion. Note that these BINs could be present for others species within the GSL-rl.**

| Species | BIN | Reason of exclusion |
|---|---|---|
| *Acanthodoris pilosa* | BOLD:ACO2942 | Another BIN is more geographically relevant |
| *Acanthodoris pilosa* | BOLD:AAX0711 | Another BIN is more geographically relevant |
| *Anarhichas denticulatus* | BOLD:ACE4267 | Highly potential wrong identification (*Anarchinas minor*) |
| *Arctogadus glacialis* | BOLD:AAA9472 | Highly potential wrong identification (*Boreogadus saida*) |
| *Artediellus atlanticus* | BOLD:AAB1020 | Highly potential wrong identification (*Triglops murrayi*) |
| *Aspidophoroides monopterygius* | BOLD:AAA9929 | Highly potential wrong identification |
| *Balanus crenatus* | BOLD:AEB7035 | Another BIN is more geographically relevant |
| *Balanus crenatus* | BOLD:AAB1410 | Highly potential wrong identification |
| *Chaetozone setosa* | BOLD:AAB5820 | Another BIN is more geographically relevant |
| *Cirratulus cirratus* | BOLD:AAJ0882 | Another BIN is more geographically relevant |
| *Cirratulus cirratus* | BOLD:ACJ5831 | Another BIN is more geographically relevant |
| *Cirratulus cirratus* | BOLD:ACH1141 | Another BIN is more geographically relevant |
| *Coryphella verrucosa* | BOLD:AAN6579 | Highly potential wrong identification |
| *Cyanea capillata* | BOLD:AAF9673 | Another BIN is more geographically relevant |
| *Cyanea capillata* | BOLD:ACM6954 | Another BIN is more geographically relevant |
| *Doto coronata* | BOLD:AAE8508 | Another BIN is more geographically relevant |
| *Doto coronata* | BOLD:ACB8287 | Another BIN is more geographically relevant |
| *Eualus pusiolus* | BOLD:ABW0161 | Another BIN is more geographically relevant |
| *Laonice cirrata* | BOLD:ACH1038 | Another BIN is more geographically relevant |
| *Laonice cirrata* | BOLD:ACC2851 | Another BIN is more geographically relevant |
| *Laonice cirrata* | BOLD:ACI3025 | Another BIN is more geographically relevant |
| *Laonice cirrata* | BOLD:AAV0397 | Another BIN is more geographically relevant |
| *Laonice cirrata* | BOLD:ACV6427 | Another BIN is more geographically relevant |
| *Limnodrilus hoffmeisteri* | BOLD:ADK1554 | Another BIN is more geographically relevant |
| *Limnodrilus hoffmeisteri* | BOLD:ACW0660 | Another BIN is more geographically relevant |
| *Littorina saxatilis* | BOLD:AAA7469 | Highly potential wrong identification |
| *Micrura varicolor* | BOLD:AAK8914 | Highly potential wrong identification |
| *Obelia dichotoma* | BOLD:AAA7089 | Highly potential wrong identification |
| *Obelia geniculata* | BOLD:AAA7088 | Another BIN is more geographically relevant |
| *Obelia geniculata* | BOLD:ADK7351 | Another BIN is more geographically relevant |
| *Obelia longissima* | BOLD:AAF4295 | Highly potential wrong identification |
| *Oithona similis* | BOLD:AAD7750 | Another BIN is more geographically relevant |
| *Oithona similis* | BOLD:ACQ4233 | Another BIN is more geographically relevant |
| *Onchidoris muricata* | BOLD:AAZ3494 | Highly potential wrong identification |
| *Orthopyxis integra* | BOLD:AAE5348 | Highly potential wrong identification |
| *Orthopyxis integra* | BOLD:AAE5349 | Highly potential wrong identification |
| *Orthopyxis integra* | BOLD:AAE5350 | Highly potential wrong identification |
| *Orthopyxis integra* | BOLD:AAE5351 | Highly potential wrong identification |
| *Pandalus montagui* | BOLD:AAB2200 | Highly potential wrong identification |

| | | |
|---|---|---|
| *Phycis chesteri* | BOLD:AAB0057 | Highly potential wrong identification (*Merluccius bilinearis*) |
| *Polymastia boletiformis* | BOLD:ACQ0787 | Highly potential wrong identification |
| *Praxillella affinis* | BOLD:ACH1056 | Another BIN is more geographically relevant |
| *Praxillella praetermissa* | BOLD:ACH1056 | Another BIN is more geographically relevant |
| *Spiophanes bombyx* | BOLD:ACH1173 | Another BIN is more geographically relevant |
| *Spiophanes bombyx* | BOLD:ACH1123 | Another BIN is more geographically relevant |
| *Spirorbis spirorbis* | BOLD:AAJ3464 | Highly potential wrong identification |
| *Stomias boa* | BOLD:AAB5302 | Another BIN is more geographically relevant |
| *Temora stylifera* | BOLD:ADK4239 | Another BIN is more geographically relevant |
| *Temora stylifera* | BOLD:ADK0428 | Another BIN is more geographically relevant |
| *Triglops nybelini* | BOLD:AAB1020 | Highly potential wrong identification |
| *Triglops nybelini* | BOLD:ADQ8959 | Highly potential wrong identification |
| *Trochochaeta multisetosa* | BOLD:ADH7281 | Another BIN is more geographically relevant |
| *Tubifex tubifex* | BOLD:AEB6122 | Another BIN is more geographically relevant |
| *Tubifex tubifex* | BOLD:AAA6805 | Another BIN is more geographically relevant |
| *Urasterias lincki* | BOLD:AAG0209 | Highly potential wrong identification |
| *Weberella bursa* | BOLD:ACH5414 | Highly potential wrong identification |

### C. Creation of an eDNA metabarcoding dataset

Water samples of 2L were collected from scientific surveys in 2018 in coastal areas from the GSL, both at surface and bottom of the water column (n=61, Fig. S2). Negative field controls (n=2) were also obtained by pouring 2L of clean water at sampling site. Water samples and controls were then frozen at -20$^{o}$C until their transport to Maurice Lamontagne Institute and afterwards at -40$^{o}$C until filtration. Water samples and controls were thawed and filtered on glass fiber 47 mm, 1.2 and/or 10 µm pore size in an ultraclean room (n = 63 filters) and 10 negative filtration controls were added. Qiagen Blood and Tissus Kit was used for DNA extraction from filters, including 7 extraction controls (elution in 80µl AE buffer). PCR were performed using 5  µl of a 1/10 dilution of the extract at Genome Québec with the primers mICOIintF (Leray et al. 2013) and jgHCO2198 (Geller et al. 2013), targeting a 313 pb section of the COI Folmer region. Three PCR negative controls were added for a total of 83 samples. Each amplification was then indexed and multiplexed, and then sequenced using Illumina HiSeq 6000 PE 250 pb at Genome Québec.

Reads were demultiplexed by the sequencing facility, and quality control of raw reads was assessed using FastQC (Andrews 2010) and MultiQC (Ewels et al. 2016). Adapters were retrieved and removed using cutadapt (Martin 2011). Then, the dada2 R package (Callahan et al. 2016) was used to create exact sequence variants (ESV) from the data. Briefly, the *trimandfilter* function was first used for quality filtering with slightly modified parameters (truncQ = 10, maxEE = 2;), then error rate was computed with the *learnErrors* function. Reads were then dereplicated (*derepFastq* function), samples were inferred (*dada* function) then paired reads were merged (*mergePairs* function, minOverlap = 30, maxMismatch = 0). An ESV table was created (*makeSequenceTable* function), with samples as columns and ESV as rows, and chimeras were removed with the *removeBimeraDenovo* function. The ESV table was further corrected based on negative samples, where we have subtracted from the total number of observations the number observed in related negative controls (from sampling to PCR control). ESVs with only one read detected were removed. More details on the bioinformatics pipeline can be find on Fig. S3.

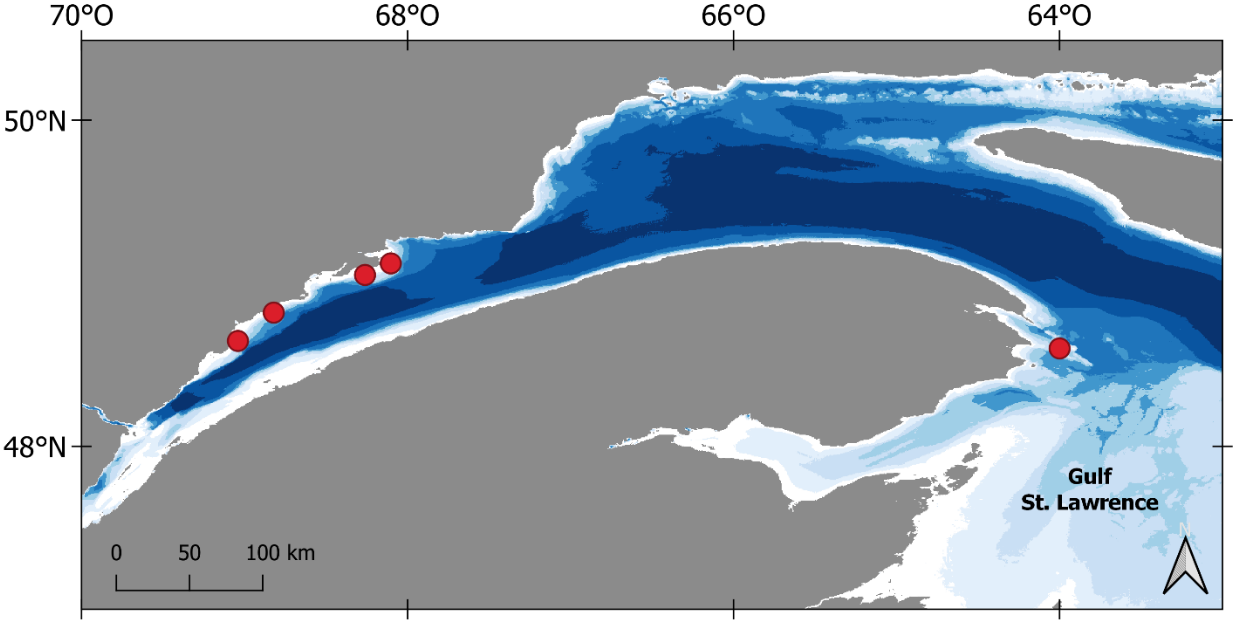

**Figure S2 Sampling locations for the metabarcoding dataset in the St. Lawrence.**

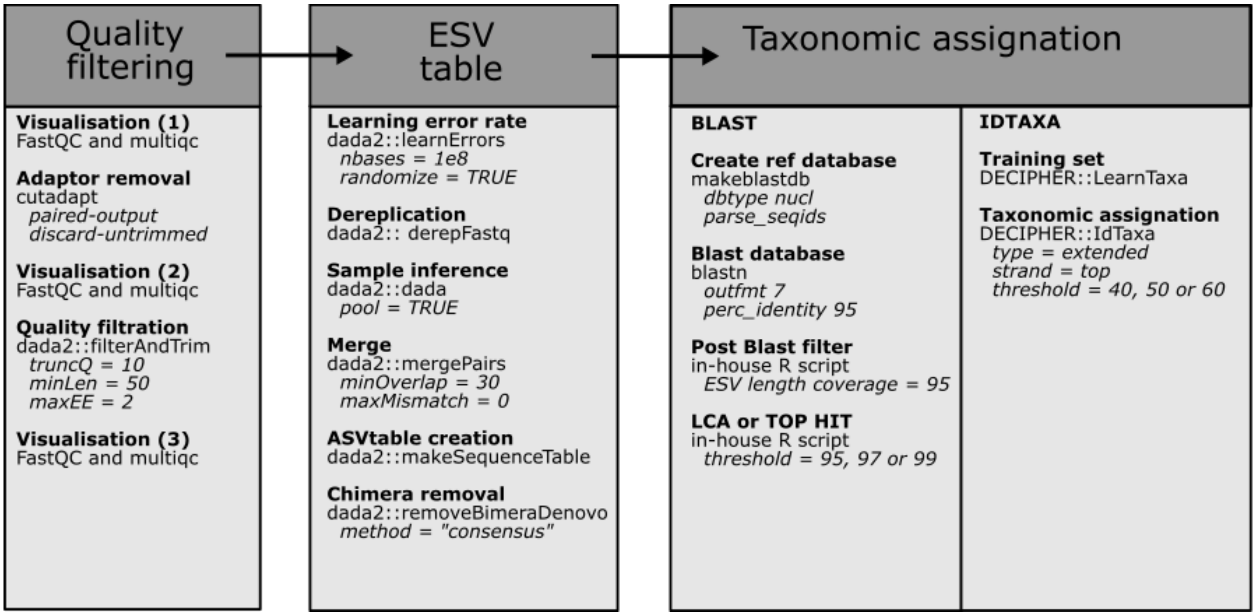


**Figure S3 Schematic representation of the metabarcoding bioinformatics pipeline.**
